# eIF2B localisation and its regulation during the integrated stress response is cell type specific

**DOI:** 10.1101/2023.01.10.523470

**Authors:** Filipe M. Hanson, Madalena I. Ribeiro de Oliveira, Alison K. Cross, K. Elizabeth Allen, Susan G. Campbell

## Abstract

Eukaryotic initiation factor 2B (eIF2B) is a master regulator of translation control. eIF2B recycles inactive eIF2-GDP to active eIF2-GTP. Under transient/acute cellular stress, a family of kinases phosphorylate the alpha subunit of eIF2 (eIF2α-P[S51]) activating the integrated stress response (ISR). This response pathway inhibits eIF2B activity resulting in overall translation attenuation and reprogramming of gene expression to overcome the stress. The duration of an ISR programme can dictate cell fate wherein chronic activation has pathological outcomes. Vanishing white matter disease (VWMD) is a chronic ISR-related disorder linked to mutations in eIF2B. eIF2B is vital to all cell types, yet VWMD eIF2B mutations primarily affect astrocytes and oligodendrocytes suggesting cell type-specific functions of eIF2B. Regulation of the cytoplasmic localisation of eIF2B (eIF2B bodies) has been implicated in the ISR. Here, we highlight the cell type specific localisation of eIF2B within neuronal and glial cell types. Our analyses revealed that each cell type possesses its own steady-state repertoire of eIF2B bodies with varied subunit composition and activity. We also demonstrate that neural and glial cell types respond similarly to acute induction of the ISR whilst a chronic ISR programme exerts cell type-specific differences. Regulatory composition of eIF2B bodies is suggested to be differentially modulated in a manner that correlates to the action of acute and chronic ISR. We also highlight a cell type specific response of the ISR inhibitor ISRIB on eIF2B localisation and activity.

## Introduction

All biological processes are intrinsically dependent upon the highly conserved and hierarchical process of mRNA translation. A key protein complex involved in ensuring that efficient translation initiation takes place is the eukaryotic initiation factor 2, eIF2. eIF2 is a heterotrimeric G-protein made up of the subunits α, β, and γ (Naveau et al., 2013; Schmitt et al., 2012). In its active GTP-bound state, eIF2 is complexed with initiator methionyl transfer RNA (eIF2-GTP-Met-tRNAi) and forms a ternary complex (TC) whose key role is to locate the first start codon to the ribosome (Hinnebusch & Lorsch, 2012). Following codon recognition, eIF2-GTP is hydrolysed to eIF2-GDP through the action of the canonical GTPase-activating protein eIF5 (Paulin et al., 2001). Crucial for successive rounds of translation is the regeneration of GTP-bound eIF2 which is catalysed by the guanine nucleotide exchange factor (GEF) eIF2B. Once released from the scanning ribosome, eIF5 stays associated with eIF2-GDP and hinders any spontaneous GDP release (GDP dissociation inhibitor, GDI) from eIF2. In addition to its GEF function, eIF2B acts as a GDI displacement factor (Jennings et al., 2013), removing eIF5, followed by GDP release from eIF2 (Williams et al., 2001). These functions highlight eIF2B as a powerful control checkpoint for the availability of TCs.

In its native form, eIF2B is a heterodecameric complex composed of two copies of 5 non-identical subunits (termed eIF2Bα-ε). The γ and ε subunits catalyse the GEF activity, whereas the α, β and δ subunits regulate this activity in response to different cellular stress insults (Bogorad et al., 2014; Kimball et al., 1998; Pavitt et al., 1997; Pavitt et al., 1998). Structurally, eIF2B decameric conformation is comprised of an eIF2B(αβδ)_2_ hexameric regulatory core laid between two eIF2B(γε) catalytic heterodimers (Tsai et al., 2018; Zyryanova et al., 2018). In mammalian cells, eIF2B has been reported to exist in different sub-complexes arrangements with varying subunit composition (Liu, et al., 2011; Wortham et al., 2014).

At the hub of translational control is the regulation of eIF2B activity by the integrated stress response (ISR) (Pakos-Zebrucka et al., 2016; Hanson et al., 2022). During acute or transient stress, the ISR activates stress-sensing kinases (PERK, PKR, GCN2, HRI) which phosphorylate the α subunit of eIF2 at serine 51 (eIF2α-P[S51]). Phosphorylated eIF2α acts as a competitive substrate to its unphosphorylated cognate, blocking GEF activity of decameric eIF2B by inhibiting the interaction of eIF2γ with the eIF2Bε subunit (Schoof et al., 2021; Zyryanova et al., 2021; Adomavicius et al., 2019; Kashiwagi et al., 2017; Kashiwagi et al., 2019; Kashiwagi et al., 2016). Attenuated eIF2B activity limits TC levels and reduces global protein synthesis. Concomitantly, a specific subset of mRNAs harbouring upstream ORFs bypass this translation attenuation. These include activating transcription factor 4, ATF4, and C/EBP homologous protein, CHOP (Harding et al., 2000). In contrast, transition to a chronically activated ISR is widely reported as adaptive to prolonged stress, ultimately pro-apoptotic when cells are unable to overcome sustained stress with pathological consequences (Bond et al., 2020).

In yeast cells, eIF2B localises to stable cytoplasmic foci termed ‘eIF2B bodies’ where GEF activity takes place and are targeted for regulation (Campbell et al., 2005; Campbell & Ashe, 2006; Egbe et al., 2015; Moon & Parker, 2018; Norris et al., 2021; Nüske et al., 2020; Taylor et al., 2010). These studies were further extended in mammalian cells where heterogeneous populations of different-sized bodies correlating to their eIF2B subunit makeup were observed (Hodgson et al., 2019). Larger bodies contained all eIF2B subunits, whilst small bodies predominantly consisted of the γ and ε catalytic subunits. Upon acute endoplasmic reticulum (ER) stress, it was demonstrated that the ISR differentially modulates these eIF2B body subpopulations, decreasing the GEF activity of larger bodies and inversely increasing GEF activity within small bodies. This increase in GEF activity was concomitant with a redistribution of eIF2Bδ to small bodies, suggesting the existence of a previously unidentified eIF2Bγδε heterotrimeric sub-complex. ISR-targeting drugs (*e.g*. ISRIB) which boost translation recapitulated this eIF2Bδ redistribution to small bodies in unstressed cells (Hodgson et al., 2019), thus implying that this action might be an innate response to the ISR to allow low baseline levels of translation. Nonetheless, the functional relevance of eIF2Bδ redistribution is still unknown.

Despite eIF2B’s ubiquitous role in the ISR across all cell types (Pakos-Zebrucka et al., 2016), mutations in any of the five subunits of eIF2B result in the neurological disorder leukodystrophy with vanishing white matter disease (VWMD) (van der Knaap et al., 2006). VWMD mutations are selectively detrimental to astrocytes, cause defective maturation and mitochondrial dysfunction in oligodendrocytes and, ultimately, lead to neuronal death due to axonal de-myelination (Bugiani et al., 2011; Dooves et al., 2016; Dooves et al., 2018; Herrero et al., 2019; Klok et al., 2018; Leferink et al., 2018). Surprisingly, studies have shown that cultured neurons are unaffected by eIF2B VWMD mutations, implying that cell type-specific features of eIF2B function and regulation may exist at least in brain cell types, which remains to be understood. We previously showed that eIF2B bodies are sites of eIF2B GEF activity as eIF2 can shuttle into these bodies in a manner that correlates with ISR activation (Hodgson et al., 2019). Here, we investigated steady-state eIF2B localisation dynamics and subsequent changes upon cellular stress and classical ISR-targeting drugs in neuronal and glial cell lines. We report that eIF2B localisation to eIF2B bodies is tailored in a cell type-specific manner. We also demonstrate that the regulatory composition of eIF2B bodies is tightly modulated by cellular stress in a cell type-manner. We further showcase a novel cell type-sensitivity feature of ISRIB in the regulation of eIF2B body composition and eIF2 shuttling.

## Results

### eIF2B localises to eIF2B bodies in a cell type dependent manner

eIF2B localisation has been reported in yeast (Campbell et al., 2005; Moon & Parker, 2018; Taylor et al., 2010) and, more recently, in mammalian cells (Hodgson et al., 2019), however the latter shows a higher degree of complexity. To further our knowledge of cellular eIF2B localisation, we transiently transfected the catalytic ε subunit (eIF2Bε) tagged with a monomeric green fluorescent protein (mGFP) into neuroblastoma (SH-SY5Y), astrocytoma (U373) and hybrid primary oligodendrocytes (MO3.13) cell lines and observed different patterns of eIF2B localisation in all 3 cells lines (Figure 1A and Figure S1A). Cells expressing eIF2Bε-mGFP exhibited either eIF2B bodies or the localisation was fully dispersed throughout the cytoplasm (Figure 1Bi). We observed that the percentage (%) cells localising eIF2B significantly differs across cell types (Figure 1Bii). U373 cells showed the highest percentage of cells containing eIF2B bodies (53.50%) followed by MO3.13 (33.25%) and SH-SY5Y exhibiting the lowest percentage (19.25%). Because eIF2B overexpression could potentially impact the observed localisation pattern across cell types, we examined endogenous eIF2Bε and observed a similar trend (Figure S1B). Next, given the heterogenous populations of different sized eIF2B bodies, we subcategorised them into small eIF2B bodies (<1μ^2^) or large eIF2B bodies (≥1μ^2^) (Figure 1Ci). Small eIF2Bε-mGFP bodies were the predominant subpopulation across all cell types. U373 and MO3.13 cells exhibited a similar percentage per cell (88.19% and 89.34%, respectively), and both were slightly higher in comparison to SH-SY5Y cells (71.46%). In contrast, SH-SY5Y cells displayed an increased average percentage of large eIF2Bε-mGFP bodies per cell (30.54%) in comparison to U373 and MO3.13 cells (13.81% and 12.66%, respectively). Here, we show that eIF2B localisation is fundamentally cell type specific: each brain cell type harbours its own prevalence of eIF2B bodies although abundance of each body size group is suggested to be similar amongst glial cell types.

**Figure 1.**
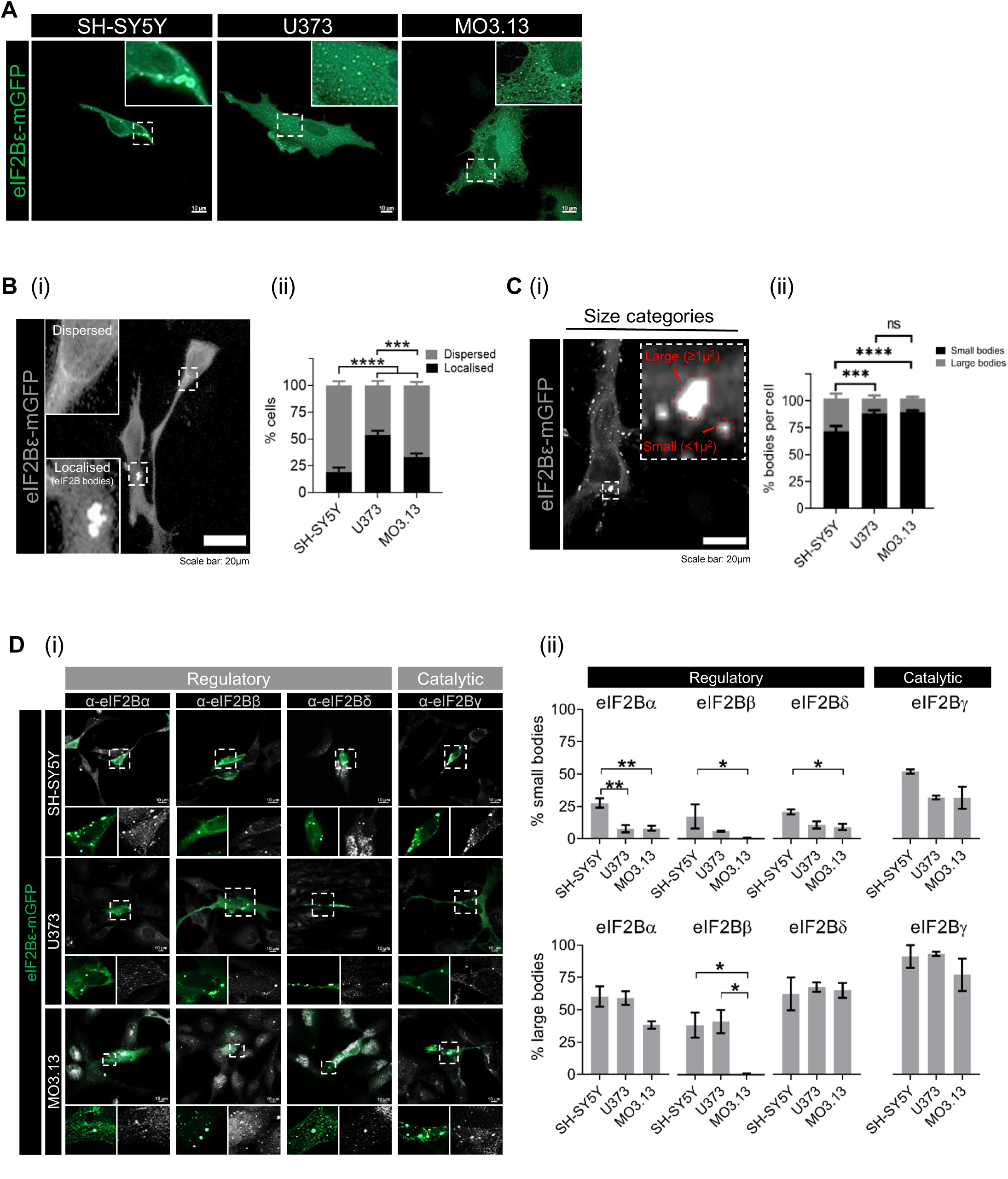
eIF2B localisation is cell type-specific. **(A)** SH-SY5Y, U373 and MO3.13 cells subjected to transient transfection and expressing eIF2Bε-mGFP. Scale bar: 10 μm. **(B) (i)** Cells express dispersed eIF2B or localised eIF2B (eIF2B bodies). **(ii)** Mean percentage of cells displaying dispersed eIF2B or localised eIF2B in a population of 100 transfected cells (mean±SEM; *N*=4; \*\*\*\**p*≤0.001, \*\*\**p*≤0.001 according to two-way ANOVA). **(iii)** eIF2B bodies were categorised as small bodies (<1μ^2^) and large bodies (≥1μ^2^). **(ii)** Within the transfected cells exhibiting localised eIF2B, the mean percentage of small and large eIF2B bodies in a population of 50 cells (mean±SEM; *N*=3; \*\*\*\**p*≤0.001, \*\*\**p*≤0.001, *ns*: non-significant according to two-way ANOVA). **(D) (i)** Confocal images of SH-SY5Y, U373 and MO3.13 expressing eIF2Bε-mGFP and immunolabelled with primary anti-eIF2Bα, anti-eIF2Bβ, anti-eIF2Bδ and anti-eIF2Bγ. Scale bar: 10 μm. **(ii)** Mean percentage of transfected cells displaying co-localisation of eIF2B(α-γ) subunits to small (*top panel*) and large (*bottom panel*) eIF2Bε-mGFP bodies of at least 30 cells per repeat (mean±SEM; *N*=3; \*\**p*≤0.01, \**p*<0.05 according to one-way ANOVA).

### Subunit composition of eIF2B bodies is cell type-specific

eIF2B exists as a decameric complex. eIF2Bε alone can carry out GEF activity, however the rate of this exchange is enhanced upon joining of other eIF2B subunits (Gomez & Pavitt, 2000). Regulatory subunits increase GEF activity, modulate levels of eIF2B activity upon cellular stress and, more recently, colocalise to eIF2B bodies in a size-dependent manner (Liu et al., 2011; Hodgson et al., 2019). Having shown eIF2B localisation is different between cell types (**Figure 1B and Figure 1C**), we next investigated whether subunit co-localisation to eIF2Bε-mGFP bodies also exhibits cell type specific features. We performed immunocytochemistry on the regulatory (eIF2Bα, eIF2Bβ, eIF2Bδ) and catalytic (eIF2Bγ) subunits of eIF2B in SH-SY5Y, U373 and MO3.13 cells (**Figure 1Di**). Previous data using U373 cells revealed that small eIF2B bodies predominantly contain catalytic subunits, whilst larger eIF2B bodies additionally contain a mixture of regulatory subunits (Hodgson et al., 2019). We confirmed that this trend is observed across all cell types by measuring the percentage (%) of small and large eIF2Bε-mGFP bodies that co-localise with the remaining subunits (eIF2Bα-γ). eIF2Bγ co-localisation with eIF2Bε-mGFP showed the highest mean percentage in small eIF2B bodies, although slightly increased in neuronal cells when compared to glial cells (SH-SY5Y: 51.99%; U373: 31.86%; MO3.13: 31.63%) (**Figure 1Dii**). Moreover, neuronal cells also displayed a significantly higher percentage of small bodies containing regulatory subunits eIF2Bα (SH-SY5Y: 27.58%; U373: 7.72%; MO3.13: 8.13%), eIF2Bβ (SH-SY5Y: 17.33%; U373: 5.94%; MO3.13: 0.68%) and eIF2Bδ (SH-SY5Y: 20.83%; U373: 10.63%; MO3.13: 9.03%). Large eIF2B bodies showed similar catalytic eIF2Bγ co-localisation across all cell types (SH-SY5Y: 91.23%; U373: 93.22%; MO3.13: 77.02%) with drastic cell-type disparities on regulatory subunit make-up (**Figure 1Dii**). Oligodendrocytic cells displayed slightly lower eIF2Bα co-localisation albeit no statistically significant difference compared to the other cell types (SH-SY5Y: 60.26%; U373: 59.02%; MO3.13: 38.25%) and near absence of eIF2Bβ co-localisation (SH-SY5Y: 38.38%; U373: 41.13%; MO3.13: 0.62%) even though endogenous eIF2Bβ localises to cytoplasmic foci (**Figure 1Dii).** eIF2Bδ co-localisation to large eIF2B bodies was overall similar across cell types (SH-SY5Y: 62.39%; U373: 67.48%; MO3.13: 65.00%). These results demonstrate that our previous findings correlating eIF2B body size to subunit composition (Hodgson et al., 2019) is somewhat exerted on a cell type basis: astrocytic and neuronal cells follow this size:subunit pattern whilst eIF2B bodies of oligodendrocytes are largely depleted of a regulatory eIF2B subunit.

### The rate of eIF2 shuttling within eIF2B bodies is cell type specific

eIF2B controls the levels of ternary complexes by regulating the available pool of GTP-bound eIF2. Previous studies have shown that the shuttling rate of eIF2 through eIF2B bodies can infer eIF2B GEF activity (Campbell et al., 2005; Campbell & Ashe, 2006; Hodgson et al., 2019; Norris et al., 2021). We co-transfected eIF2α-tGFP <u>i</u>and eIF2Bε-mRFP in SH-SY5Y, U373 and MO3.13 cells and performed fluorescence recovery after photobleaching (FRAP) on small and large eIF2B bodies. We first confirmed that all sized eIF2Bε-RFP bodies co-localised with eIF2α-tGFP (**Figure 2A**). Next, FRAP analysis showed that eIF2α-tGFP recovery of small eIF2B bodies was relatively similar across cell types, although slightly higher for U373 cells despite not being statistically significant (SH-SY5Y, 34.21%; U373, 42.32%; MO3.13, 34.16%) (**Figure 2Bi and ii**). Surprisingly, large eIF2B bodies showed drastic discrepancies. SH-SY5Y and U373 cells exhibited similar eIF2α-tGFP recovery (SH-SY5Y: 36.13%; U373: 37.08%) whilst MO3.13 cells have significantly lower recovery (22.51%) (**Figure 2Bii**). Hence. these data demonstrate that small eIF2B bodies displaying similar % recoveries are functionally similar across all cell types whilst large eIF2B bodies display cell type specific differences.

**Figure 2.**
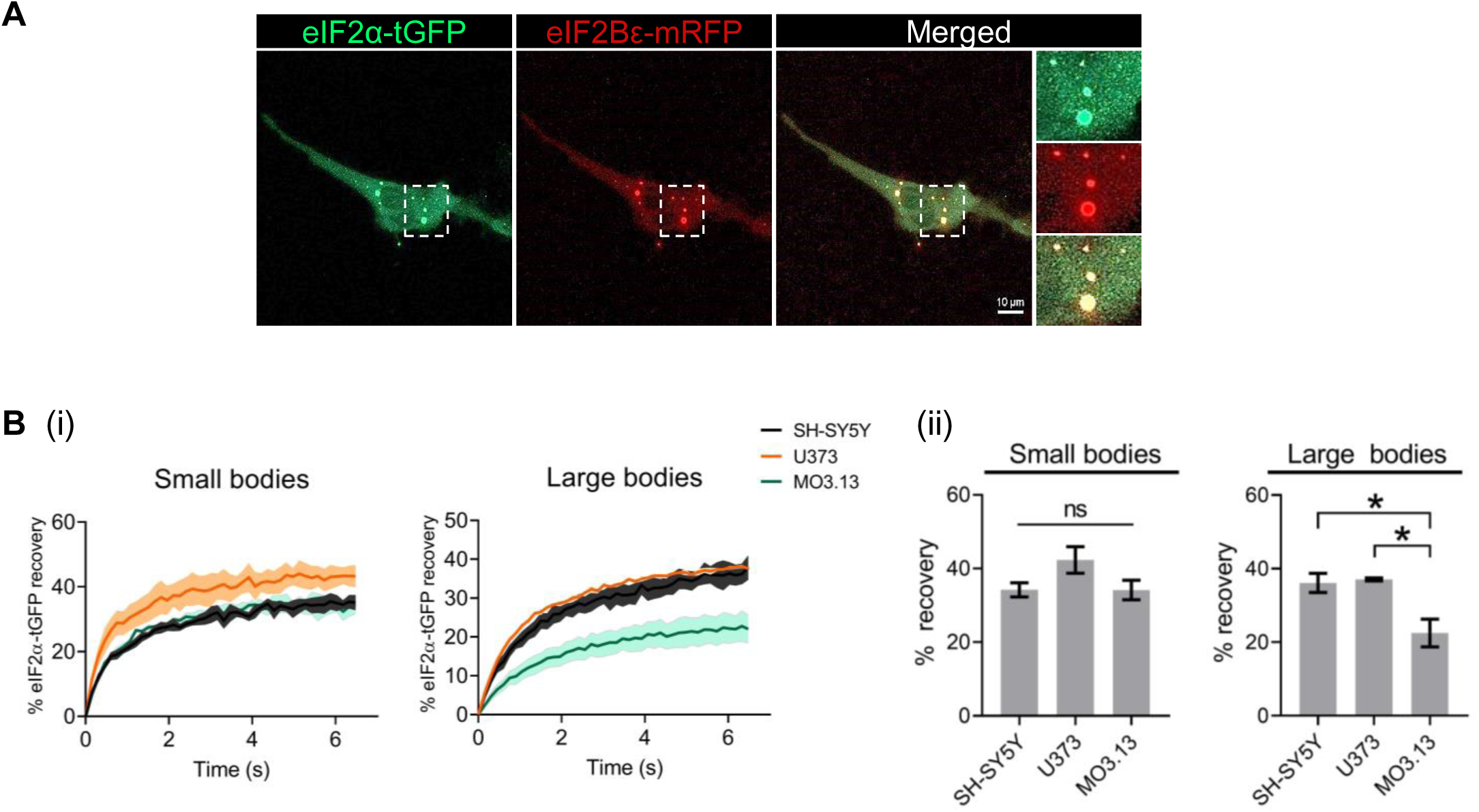
eIF2 shuttling of eIF2B bodies is cell type-specific. Cells were co-transfected with eIF2α-tGFP to carry out fluorescence recovery after photobleaching (FRAP) analysis, and eIF2Bε-mRFP to locate the eIF2B body. **(A)** Representative live cell imaging of a cell co-expressing eIF2α-tGFP and eIF2Bε-RFP. Scale bar: 10 μm. **(B) (i)** Quantification of normalised FRAP curves for eIF2α-tGFP of at least 10 bodies of each size category of SH-SY5Y, U373 and MO3.13 cells. The data were graphed and shown as the mean and SEM bands (*N*=*3*). **(ii)** Mean percentage of eIF2α-tGFP recovery determined from normalised FRAP curves (mean±SEM; *N*=3; \**p*<0.05 according to one-way ANOVA).

### The acute ISR is similar across cell types while chronic ISR displays cell type specific features

eIF2B localisation is modulated upon induction of an acute ISR in astrocytes (Hodgson et al., 2019). Here we further characterised eIF2B localisation during the transition to a chronic ISR by firstly characterising the acute *vs* chronic ISR activation in neuronal and glial cell types. To test induction of the acute ISR we used thapsigargin (Tg) and sodium arsenite (SA) to trigger ER stress and oxidative stress, respectively (**Figure 3A**). We performed western blot analysis using canonical ISR markers (PERK-P, eIF2α-P, CHOP and GADD34) (**Figure 3Bi**). As expected, short-term treatment with either Tg (1μM 1h) or SA (125μM 0.5h) led to increased eIF2α phosphorylation (eIF2α-P) and eIF2α-P-dependent protein synthesis shutdown across all cell types (**Figure 3Bii**). Next, cells were exposed to Tg at a lower concentration (300 nM) for 24h to monitor ER stress during the chronic ISR adaptation phase (Smith et al., 2020) (**Figure 3A**). As expected, PERK remained partially phosphorylated (shifted PERK band) and ISR markers (CHOP, GADD34) were expressed (**Figure 3Bi**). ATF4 expression was no longer detected at the 24h time point, however temporal monitoring during this 24h period showed that it peaked 4-8h post-Tg treatment across all cell types (**Figure S2**). A Tg treatment for 24h showed partial translation recovery in comparison to 1h Tg treatment, confirming the transition to a chronic ISR program (Guan et al., 2017).

**Figure 3.**
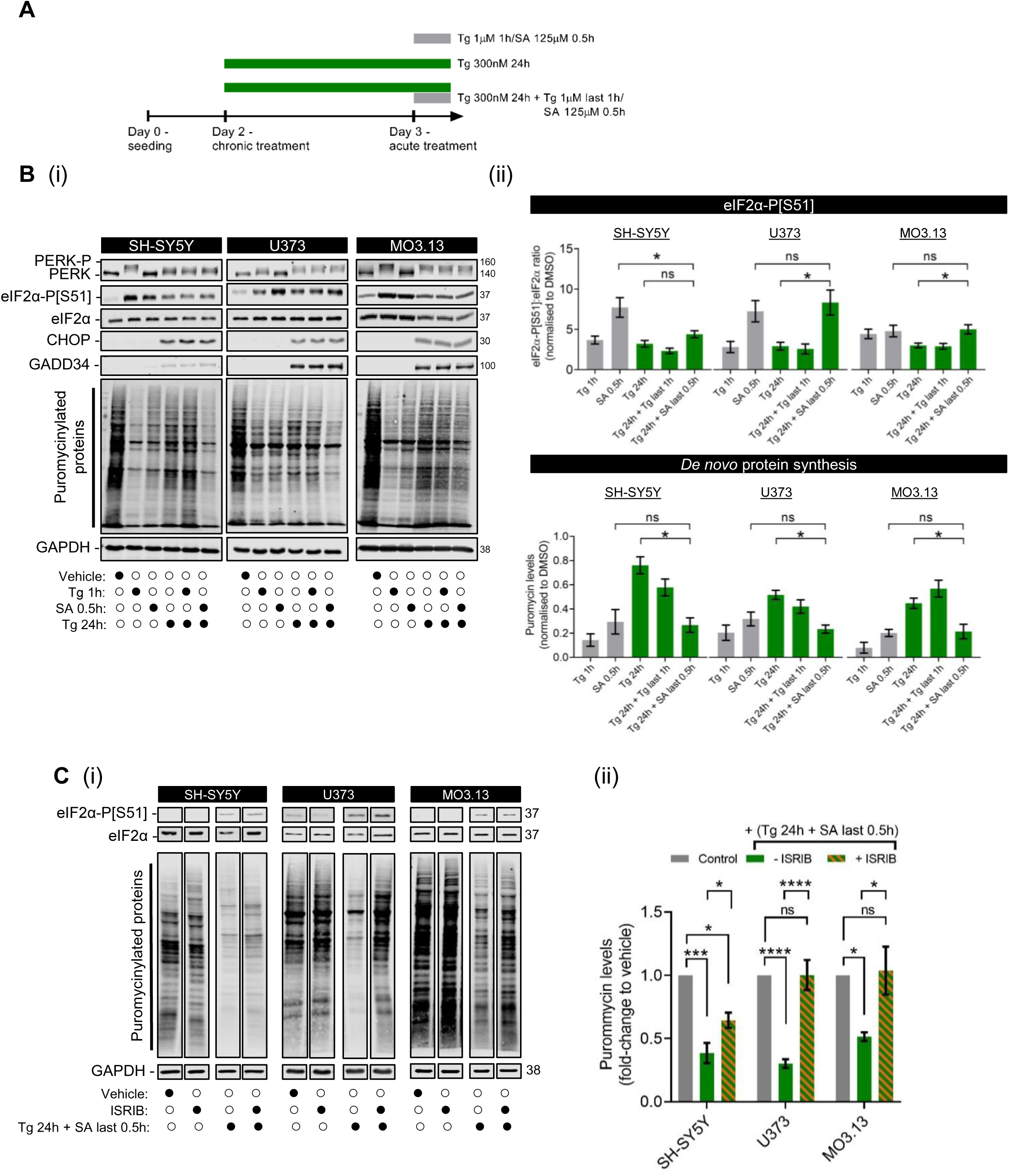
ER stress-preconditioned cells do not respond to additional acute ER stress treatment but do respond to acute oxidative stress in a cell type-manner. **(A)** Schematic diagram of stress treatments. **(B) (i)** Representative western blot of the ISR expression profile (PERK-P, PERK, eIF2α-P[S51], pan-eIF2α, CHOP and GADD34) and global newly synthesized proteins (puromycin incorporation assay) in SH-SY5Y, U373 and MO3.13 cells treated with vehicle (DMSO), acute stress inducers (Tg 1μM for 1h and SA 125μM for 30min) or chronic ER stress (Tg 300nM for 24h) subsequently challenged with previously described acute stress treatments. **(ii)** Mean expression levels of eIF2α-P[S51] normalised to total eIF2α levels (*top panel*) and puromycin-labelled nascent proteins normalised to housekeeping GAPDH levels (*bottom panel*) upon the previously described stress conditions. Fold-change relative to vehicle-treated cells was calculated and analysed using one-way ANOVA (mean±SEM; *N*=3-9; \**p*<0.05, *ns*: non-significant). Chronic ER stress conditions are highlighted in green. **(C) (i)** Representative western blot of eIF2α-P[S51], pan-eIF2α and global newly synthesized proteins (puromycin incorporation assay) in SH-SY5Y, U373 and MO3.13 cells treated with ISRIB (200nM) for 1h alone, Tg 300nM for 24h added with SA 125μM in the last 30min, or combination of both. DMSO for 24h was used as vehicle. **(ii)** Mean expression levels of puromycin-labelled nascent proteins normalised to housekeeping GAPDH levels. Fold-change relative to vehicle-treated cells was calculated and analysed using one-way ANOVA (mean±SEM; *N*=3-4; \*\*\*\**p*≤0.001, \*\*\**p*≤0.001, \**p*<0.05, *ns*: non-significant).

VWMD is predominantly characterised by an abnormal chronic-like ISR which selectively targets glial function exhibited by progressive white matter deterioration upon acute stress episodes (*e.g*. head traumas and infections). However, this glial vulnerability remains poorly understood. To provide insight into this cell type specificity, we devised a VWMD-mimicking environment in SH-SY5Y, U373 and MO3.13 cells whereby cells exposed to a chronic ISR are subsequently exposed to an acute insult. Cells were treated with 300nM Tg for 24h and then 1 μM Tg or 125 μM SA in the final 1h or 30 minutes, respectively. Interestingly, the additional Tg treatment did not affect ISR markers nor translation levels, suggesting that an on-going chronic ER stress is protective against a new ER stress insult (**Figure 3Bi and ii**). To confirm that this observed unresponsiveness was not due to Tg saturation or ISR-independent cellular effects of Tg, we treated cells with tunicamycin (Tm; which inhibits *N*-linked glycosylation of ER proteins and leads to an ER stress activated ISR like Tg) in the last 2h of a 24h treatment with 300 nM Tg. Tm treatment alone induced eIF2α phosphorylation and suppressed protein synthesis, while the additional Tm treatment to Tg preconditioned cells did not further impact protein synthesis when compared to Tg 24h alone (**Figure S3**). However, when the cells were subsequently treated with an acute oxidative stress (SA: 125 μM 0.5h), a decrease in *de novo* protein synthesis akin to SA-only levels was observed (**Figure 3Bii**), suggesting that cells reset the acute ISR program following chronic ER stress when exposed to different stressors. This decrease in protein synthesis was linked to a significant increase in eIF2α phosphorylation in U373 and MO3.13 cells. Unexpectedly, this eIF2α phosphorylation increase was not as dramatic in SH-SY5Y cells; suggesting that the suppression of protein synthesis observed here may be less dependent on eIF2α phosphorylation (**Figure 3Bii**). To test whether this was the case, we employed the same chronic stress conditions (Tg 24h, Tg 24h + Tg last 1h; Tg 24h + SA last 0.5h) in the presence or absence of the ISR inhibitor ISRIB and performed puromycin incorporation assay (**Figure 3C**). ISRIB which reverses inhibitory effects of eIF2α phosphorylation (Sidrauski et al., 2013) was unable to fully restore protein synthesis in SH-SY5Y cells compared to the glial cell types (**Figure 3C**). Taken together, these results suggest that subsequent oxidative stress in chronically ER-stressed neuronal cells is partially uncoupled from eIF2α-mediated translational control while glial cells trigger a sequential acute ISR program.

### Regulatory remodelling of small eIF2B bodies is specific to the acute phase of the ISR and partially modulated by eIF2α phosphorylation

To investigate the impact of cellular stress in eIF2B localisation, we transiently transfected SH-SY5Y, U373 and MO3.13 cells with eIF2Bε-mGFP and treated with the previously described acute (Tg 1h, SA 0.5h) and chronic (Tg 24h, Tg 24h + Tg last 1h, Tg 24h + SA last 0.5h) treatments. We observed an overall increase of eIF2B localisation in all cell types although astrocytic cells displayed a higher degree of stimulation (**Figure S4**). Furthermore, SH-SY5Y and MO3.13 cells showed significantly increased cells harbouring localised eIF2B when treated with a VWMD-devised condition (Tg 24h + SA last 0.5h) (**Figure S4**).

We previously reported increased eIF2Bδ localisation to small eIF2B bodies (mainly composed of catalytic γ and ε subunits) upon acute ISR in astrocytes, suggesting the presence of novel eIF2Bγδε subcomplexes (Hodgson et al., 2019). This implies that eIF2Bδ redistribution may play a functional role during cellular ISR, however the specific role is unknown. Given the similarities in the response to acute ISR observed in neuronal and glial cell lines (**Figure 3B**), we wanted to investigate whether the redistribution of eIF2Bδ was also similarly regulated. We performed immunofluorescence analysis using an eIF2Bδ antibody in SH-SY5Y, U373 and MO3.13 cells expressing eIF2Bε-mGFP (**Figure 4Ai**). As expected, short-term Tg and SA treatment increased eIF2Bδ localisation to small eIF2B bodies in all cell types whilst large eIF2B bodies remained predominantly unchanged (**Figure 4Aii**). During a chronic ISR treatment (Tg 24h), eIF2Bδ composition of small bodies return to control levels (**Figure 4Aii**) suggesting that this stress-induced eIF2Bδ movement is specific to an acute ISR. Surprisingly, the additional SA treatment after chronic ER stress exposure mirrored the cell type pattern observed for eIF2α phosphorylation (**Figure 3Bii**). U373 and MO3.13 cells which showed an induction of the acute ISR, also displayed a redistribution of eIF2Bδ to small eIF2B bodies resembling their respective SA-only treatment (**Figure 4Aii**). In neuronal cells, this acute SA insult after chronic ER stress, which did not induce high levels of eIF2α phosphorylation, failed to significantly enhance eIF2Bδ localisation to small eIF2B bodies when compared to levels treated with SA only (**Figure 4Aii**).

**Figure 4.**
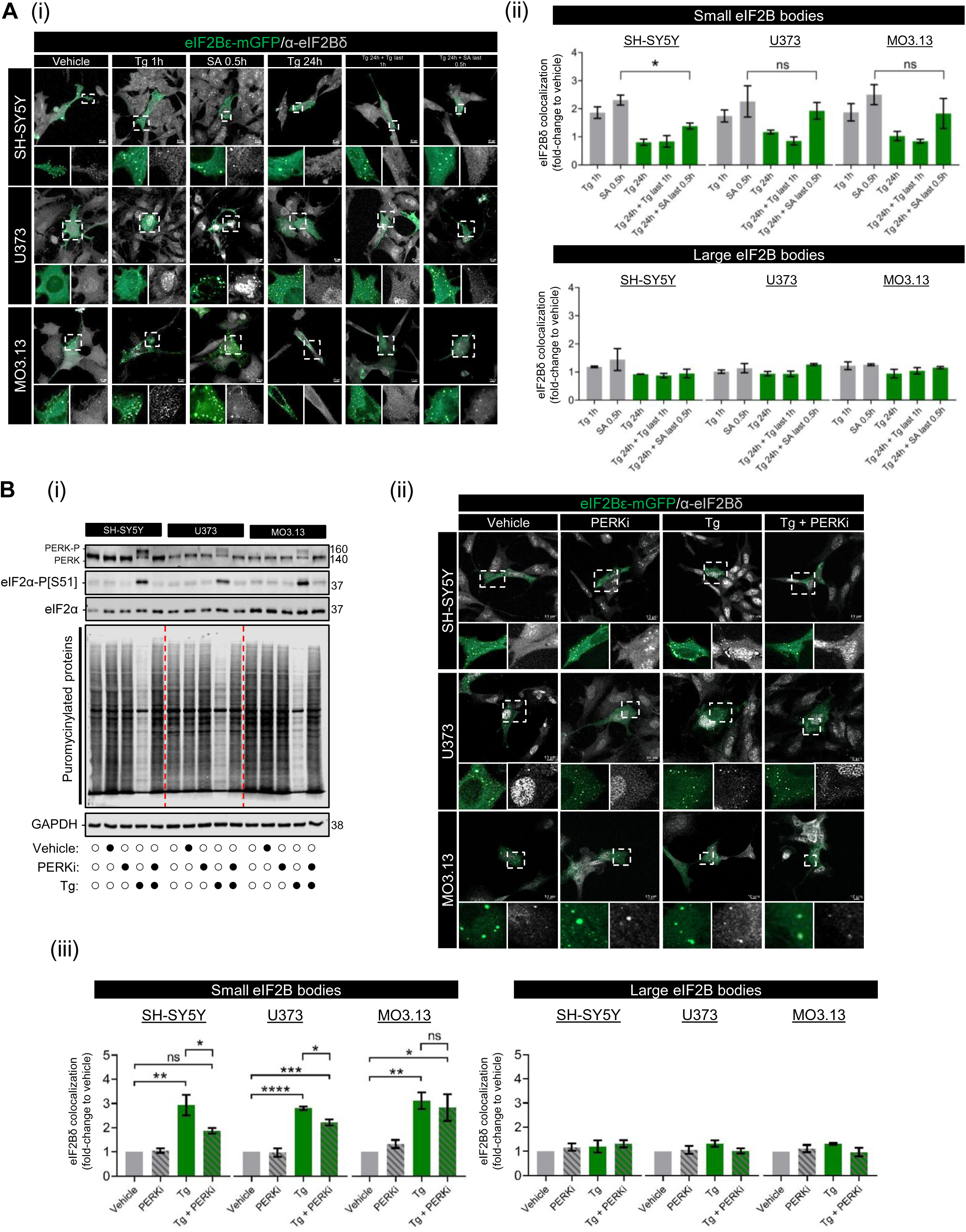
eIF2Bδ remodelling of small eIF2B bodies is transient during cellular stress and partially dictated by eIF2α-P[S51] in a cell type-dependent manner. **(A) (i)** Confocal images of SH-SY5Y, U373 and MO3.13 expressing eIF2Bε-mGFP and immunolabelled with primary anti-eIF2Bδ subjected with acute stress inducers (Tg 1μM for 1h and SA 125μM for 30min) or chronic ER stress (Tg 300nM for 24h) subsequently challenged with previously described acute stress treatments. Scale bar: 10 μm. **(ii)** Mean percentage of transfected cells displaying co-localisation of anti-eIF2Bδ to small (*top pane*l) and large (*bottom panel*) eIF2Bε-mGFP bodies of a population of 30 cells per biological repeat. Fold-change relative to vehicle-treated cells was calculated and analysed using one-way ANOVA (mean±SEM; \**p*<0.05; *ns*, non-significant). **(B) (i)** Representative western blot of the ISR expression profile (PERK-P, PERK, eIF2α-P[S51], pan-eIF2α, CHOP and GADD34), global newly synthesized proteins (puromycin incorporation assay) and loading control GAPDH in SH-SY5Y, U373 and MO3.13 cells treated with vehicle (DMSO), GSK2606414/PERKi (500 nM), Tg (1μM) or co-treated with PERKi and Tg (PERKi + Tg) for 1h. **(ii)** Confocal images of SH-SY5Y, U373 and MO3.13 cells expressing eIF2Bε-mGFP and immunolabelled with primary anti-eIF2Bδ subjected to previous treatments. Scale bar: 10 μm. **(iii)** Mean percentage of transfected cells displaying co-localisation of anti-eIF2Bδ to small (*left panel*) and large (*right panel*) eIF2Bε-mGFP bodies of a population of 30 cells per biological repeat. Fold-change relative to vehicle-treated cells was calculated and analysed using one-way ANOVA (mean±SEM; *N*=3; \**p*<0.05).

eIF2Bδ redistribution has been previously hypothesized to be modulated by levels of eIF2α-P (Hodgson et al., 2019), and here we further strengthened this hypothesis by observing a mirrored pattern of increased eIF2α-P levels and increased eIF2Bδ to small bodies (**Figure 3B and Figure 4A**). To investigate whether levels of eIF2α-P influence eIF2Bδ redistribution, we subjected cells to acute Tg treatment in the presence or absence of GSK2606414, a highly selective inhibitor of PERK (PERKi) (Halliday et al., 2017). In line with this, PERKi completely blocked eIF2α phosphorylation and inhibited translation suppression when co-treated with Tg across all cell types (**Figure 4Bi**). We again performed an immunofluorescence analysis using eIF2Bδ antibody in SH-SY5Y, U373 and MO3.13 cells expressing localised eIF2Bε-mGFP under the previously described Tg and PERKi conditions (**Figure 4Bii**). Unexpectedly, while we observed a slight increase of eIF2Bδ localisation to small bodies in SH-SY5Y and U373 cells when co-treated with Tg and PERKi (thus in the absence of eIF2α phosphorylation), it was significantly lower than when compared to Tg alone treated cells (**Figure 4Biii**). Moreover, co-treatment of PERKi and Tg treatment exhibited similar levels of eIF2Bδ in small bodies of MO3.13 cells when compared to Tg alone (**Figure 4Biii**). These data indicate that eIF2Bδ localisation to small eIF2B bodies is partially dictated by eIF2α phosphorylation in a cell type-specific manner.

### ISRIB’s mode of action on eIF2B localisation is cell type specific

ISRIB is an eIF2B activator that attenuates eIF2α-P-dependent translation suppression by promoting decamer formation and enhancing eIF2B GEF activity (Schoof et al., 2021; Zyryanova et al., 2021; Sidrauski et al., 2013). ISRIB does not impact levels of eIF2α phosphorylation *per se* but rather rescues its downstream inhibitory effect on protein synthesis. Previously, we have shown that eIF2Bδ localisation to small eIF2B bodies increased as a direct effect of ISRIB’s binding to eIF2Bδ (Hodgson et al., 2019). To test whether this is a general cellular feature, we treated SH-SY5Y, U373 and MO3.13 cells expressing eIF2Bε-mGFP with ISRIB (200 nM) for 1h and performed an immunofluorescence analysis using eIF2Bδ antibody (**Figure 5Ai**). As before, large eIF2B bodies showed no changes in eIF2Bδ composition when exposed to ISRIB alone or in combination with preconditioned Tg treatment for 24h (**Figure 5Aii**). In contrast, ISRIB treatment showed increased eIF2Bδ localisation in small bodies of U373 and MO3.13 cells while a complete absence of eIF2Bδ redistribution was observed in SH-SY5Y cells (**Figure 5Aii**). Moreover, preconditioning cells to chronic ER stress abrogated eIF2Bδ movement in MO3.13 cells upon ISRIB treatment, whereas it had no impact on U373 cells which showed eIF2Bδ redistribution in all ISRIB conditions (**Figure 5Aii**). These data provide evidence that ISRIB’s mechanism of action may involve cell type specific regulation of eIF2B localisation. Given this cell type specific impact of ISRIB in the eIF2B composition of small bodies, we next aimed to investigate if this mirrored a cell type-specific rescue of protein synthesis. Puromycin incorporation assay revealed that adding ISRIB restored protein synthesis in all cell types pre-treated with Tg for 23h (**Figure 5Bi and ii**). Taken together, ISRIB’s mode of action is suggested to not be linked to eIF2Bδ remodelling of small eIF2B bodies in neuronal and oligodendrocytic cells but may be involved in astrocytic cells.

**Figure 5.**
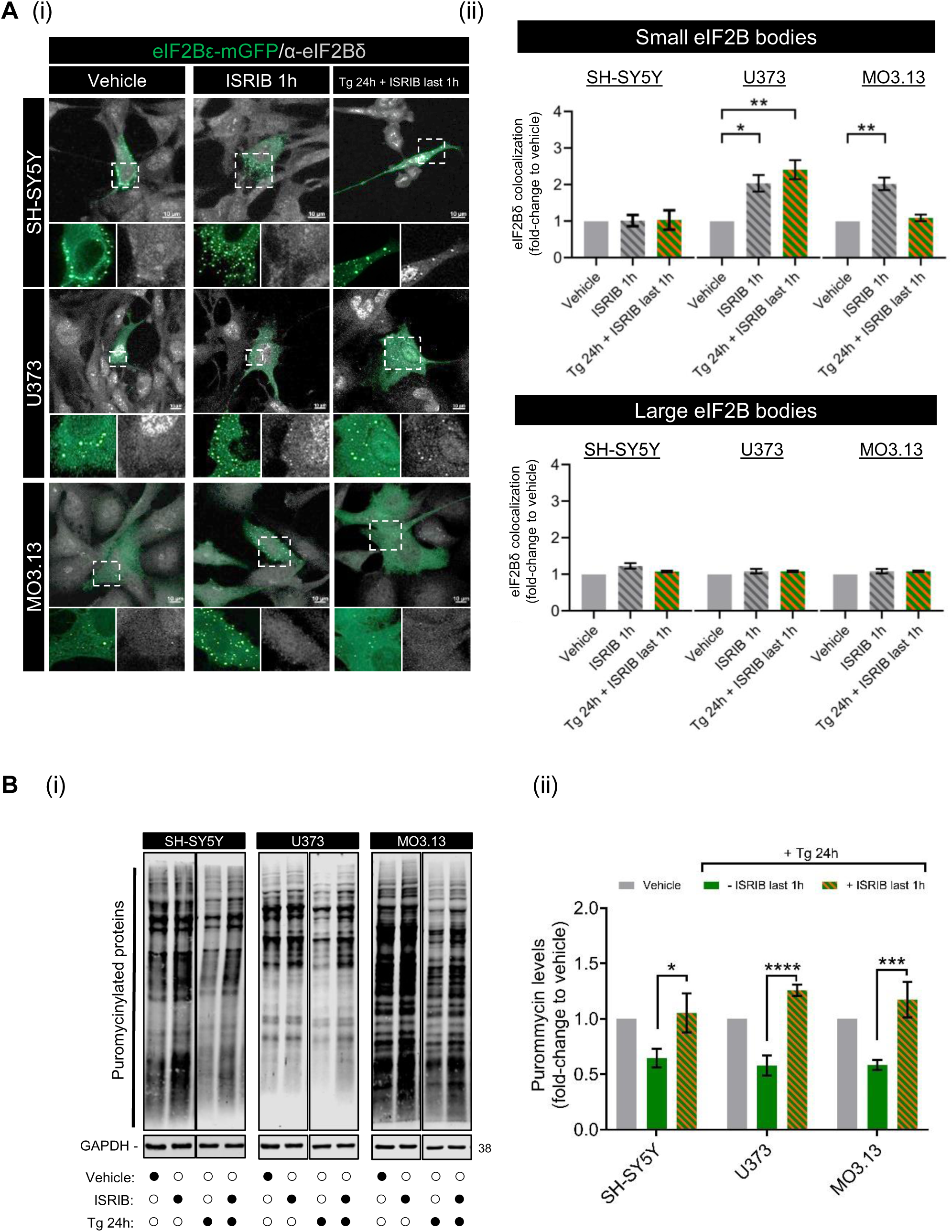
ISRIB restores translation during chronic ER stress while increasing eIF2Bδ composition of small eIF2B bodies predominantly in astrocytic cells. **(A) (i)** Confocal images of SH-SY5Y, U373 and MO3.13 expressing eIF2Bε-mGFP and immunolabelled with primary anti-eIF2Bδ subjected to ISRIB (200nM) alone 1h or in combination with preconditioned chronic ER stress treatment (Tg 300nM 24h + ISRIB last 1h). Scale bar: 10 μm. **(ii)** Mean percentage of transfected cells displaying co-localisation of anti-eIF2Bδ to small (*top panel*) and large (*bottom panel*) eIF2Bε-mGFP bodies. Fold-change relative to vehicle-treated cells was calculated and analysed using one-way ANOVA (mean±SEM; *N*=3; \*\**p*≤ 0.01, \**p*<0.05). **(C) (i)** Western blotting of global newly synthesized proteins (puromycin incorporation assay) and loading control GAPDH in SH-SY5Y, U373 and MO3.13 cells treated with same conditions as described previously (*left panel*). **(ii)** Mean expression levels of puromycin-labelled nascent proteins normalised to housekeeping GAPDH levels. Fold-change relative to vehicle-treated cells was calculated and analysed using one-way ANOVA (mean±SEM; *N*=5-9; \*\*\*\**p*≤0.001, \*\*\**p*≤0.001, \**p*<0.05).

### ISRIB and cellular stress selectively modulates activity of eIF2B bodies in a cell type-manner

In addition to the remodelling of eIF2Bδ composition in small eIF2B bodies, we have previously described that both acute stress and ISRIB result in increased shuttling of eIF2 in astrocytic cells (Hodgson et al., 2019). Therefore, we next turned to assess whether there was any cell-specific regulation of eIF2 shuttling in the different cell types upon acute and chronic cellular stress and in the presence or absence of ISRIB treatment. We treated SH-SY5Y, U373 and MO3.13 cells with ISRIB alone or with an acute Tg stress (1h) in the presence or absence of ISRIB and performed FRAP analysis on small and large eIF2B bodies. Intriguingly, cell type disparities were observed in the % recovery of eIF2 in both small and large bodies. For small bodies treated with ISRIB, a significant increase in the % recovery of eIF2 was observed for U373 cells (in line with previously published data; Hodgson et al., 2019) but not for the SH-SY5Y or MO3.13 cells (**Figure 6Ai**). Upon acute Tg stress, U373 cells displayed an increase in the % recovery of eIF2 into small bodies and this increase was sustained but not increased upon co-treatment with ISRIB (**Figure 6Ai**). Again, this increase is similar to our previous observations (Hodgson et al 2019). This increase in recovery of eIF2 in small bodies was unique to U373 cells and was not observed for either the SH-SY5Y or MO3.13 cells (**Figure 6Ai**). For large eIF2B bodies, ISRIB treatment alone did not impact on eIF2 recovery of any cell lines (**Figure 6Aii**). However, when cells were treated with an acute Tg stress (1h), a decrease in the % recovery of eIF2 was observed for both U373 and SH-SY5Y cells but not for the MO3.13 cells (**Figure 6Aii**). Furthermore, co-treatment of ISRIB and acute Tg reversed the Tg-induced inhibitory effects on eIF2 shuttling of large eIF2B bodies in U373 cells, while showing no effect on eIF2B bodies of SH-SY5Y and MO3.13 cells (**Figure 6Aii**). These data show that acute cellular stress and ISRIB predominantly regulate small and large eIF2B bodies of U373 cells amongst the cell lines used in this study.

**Figure 6.**
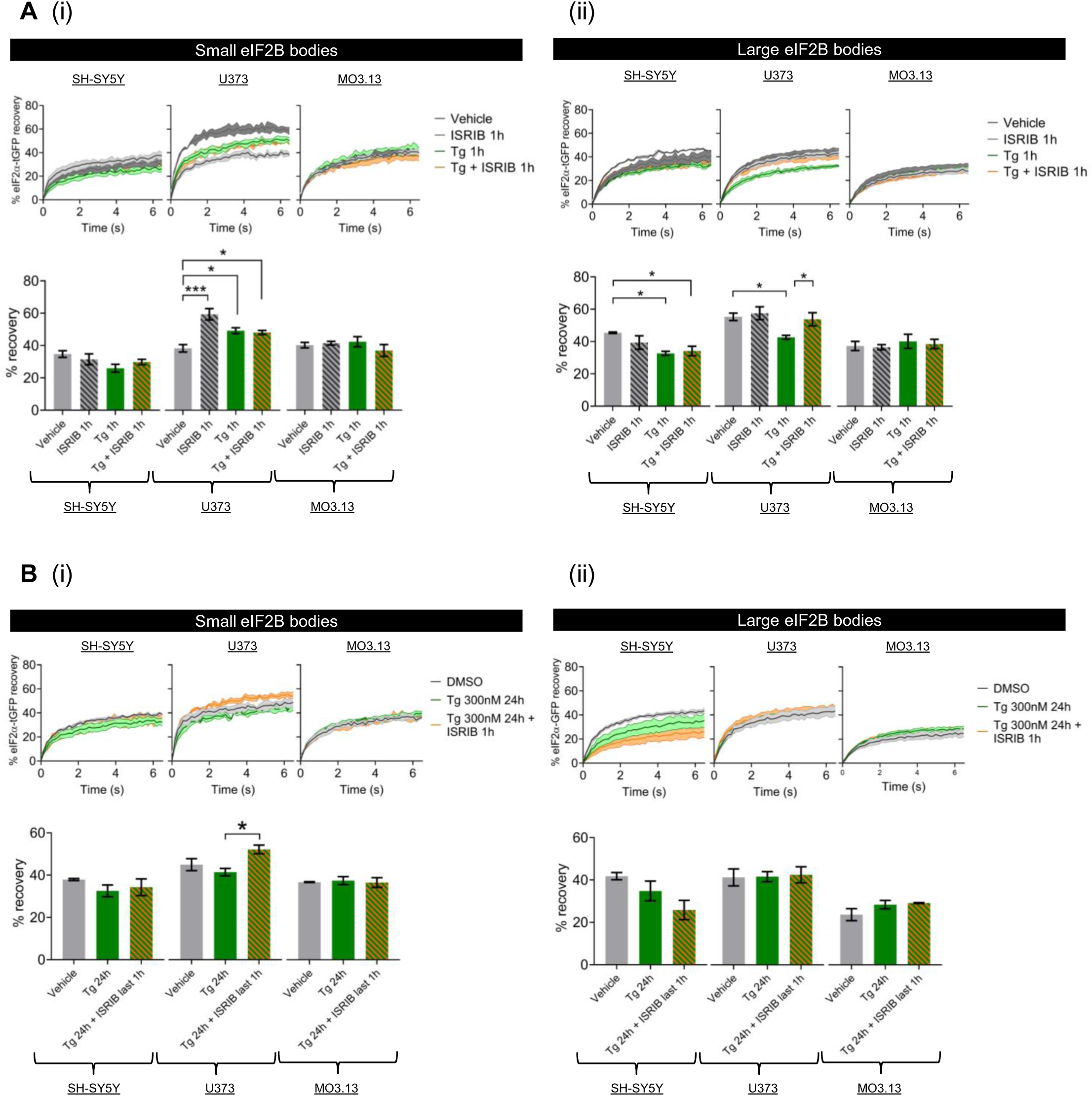
ISRIB modulates the eIF2 shuttling of eIF2B bodies in astrocytic cells. Cells were co-transfected with eIF2α-tGFP to carry out fluorescence recovery after photobleaching (FRAP) analysis, and eIF2Bε-mRFP to locate the eIF2B body. **(A)** Cells were then treated with vehicle (DMSO), ISRIB (200 nM) alone for 1h, Tg (1μM) alone for 1h or both treatments in combination (Tg + ISRIB) for 1h. Quantification of normalised FRAP curves for eIF2α-tGFP of at least 10 bodies of small (*right panel*) and large (*left panel*) eIF2Bε-mRFP bodies of SH-SY5Y, U373 and MO3.13 cells. The data were graphed and shown as the mean and S.E.M. bands (*N*=*3*). The mean percentage of eIF2α-tGFP recovery was determined from normalised FRAP curves (mean±SEM; *N*=3; \*\*\**p*≤ 0.001, \**p*<0.05 according to one-way ANOVA). **(B)** Cells were then treated with vehicle (DMSO), Tg (300nM) alone for 24h or both treatments in combination where ISRIB was added on the last hour of the 24h period exposure to Tg. Quantification of normalised FRAP curves for eIF2α-tGFP of at least 10 bodies of small (*right panel*) and large (*left panel*) eIF2Bε-mRFP bodies of SH-SY5Y, U373 and MO3.13 cells. The data were graphed and shown as the mean and S.E.M. bands (*N=3*). Mean percentage of eIF2α-tGFP recovery was determined from normalised FRAP curves (mean±SEM; *N*=3; \*\*\**p*≤ 0.001, \**p*<0.05 according to one-way ANOVA).

We next sought to observe cells with chronic ISR treatment in the presence and absence of ISRIB. In line with the recovery of protein synthesis post-chronic stress, eIF2 recovery was similar to the vehicle control cells post-24h treatment (**Figure 6Bi and ii**). Moreover, ISRIB treatment in the last hour of the 24h exposure to Tg significantly increased eIF2 recovery of small eIF2B bodies in U373 cells but not in SH-SY5Y or MO3.13 cells (**Figure 6Bi**), while altogether having no enhancing impact of the activity of large eIF2B bodies (**Figure 6Bii**). These data suggest that the activity of eIF2B bodies is transiently regulated upon cellular stress and ISRIB modulates small eIF2B bodies during chronic ISR in a cell type manner.

## Discussion

We have previously reported that eIF2B bodies represent steady-state autonomous clusters of GEF active eIF2B complexes (Campbell et al., 2005; Hodgson et al., 2019; Norris et al., 2021; Taylor et al., 2010). Here we show that the prevalence of eIF2B bodies is cell type specific in unstressed conditions (**Figure 1A-C**). Amongst the cell types used in this study, astrocytic cells showed increased number of cells displaying eIF2B bodies (~54%) in comparison to oligodendrocytic (~33%) and neuronal (~19%) cells. Because localised eIF2B accounts for only a certain portion of total eIF2B, with the remaining GEF exchange occurring elsewhere in the cytoplasm, we hypothesize that the degree of eIF2B localisation differs depending on the cellular requirement for regulated GEF activity, both for steady state purposes and ability to respond to stress.

A correlation between eIF2B body size and subunit composition was previously reported (Hodgson et al., 2019) and now we extend these studies by demonstrating that this correlation is cell type specific (**Figure 1D**). Firstly, for the small eIF2B bodies, neuronal cells harboured increased levels of regulatory subunits (eIF2Bα,β,δ) in comparison to both types of glial cells. These data indicate that small eIF2B bodies within astrocytic and oligodendrocytic cells mainly contain eIF2Bγε heterodimers, while in neuronal cells these small bodies contain eIF2B tetrameric and decameric complexes. Secondly, neuronal and astrocytic cells followed the size:subunit relationship as all four subunits (eIF2Bα-γ) showed a higher degree of co-localisation to large bodies compared to the small bodies; while oligodendrocytes exhibited the surprising absence of eIF2Bβ and a slightly decreased % of eIF2Bα. These results suggest the full eIF2B decameric complex reside in large eIF2B bodies of neuronal and astrocytic cells but may not in oligodendrocytic cells. Given that increased GEF activity of eIF2B correlates with the presence of regulatory eIF2B subunits (Dev et al., 2010; Fabian et al., 1997; Liu et al., 2011; Williams et al., 2001), this decreased association of eIF2Bβ with large eIF2B bodies may account for the decreased basal % eIF2 recovery observed for oligodendrocytes when compared to other cell types (**Figure 2B**).

eIF2B bodies are targeted during the acute ISR (Hodgson et al., 2019) however it remains unknown what their significance is upon transition to a chronic ISR. We first aimed to characterise the acute and chronic ISR of neuronal and glial cell lines used in this study. We report that the acute ISR is a general cellular feature as assessed by the canonical eIF2α-dependent pathway of translation shutdown in agreement to a plethora of other cell types (Guan et al., 2014; Spaan et al., 2019; Teske et al., 2011). Interestingly, we observed a cell type specific ability to reset an acute-like ISR depending on whether faced with repeated stresses or treated with a different stressor (**Figure 3B**). In all cells, an initial chronic ER stress was protective towards a second ER stress treatment. This has been shown by others where preconditioning cells to mild eIF2α phosphorylation, either through inhibition of PP1c or stress-inducing agents (Lu et al., 2004), has been shown to be cytoprotective. Strikingly, replacing the second stress with an oxidative stress elevated eIF2α phosphorylation in glial cells but not in neuronal cells, however protein synthesis was still targeted in neuronal cells suggesting that this second stress may be regulated via an eIF2α-independent mechanism. Our observations in neuronal cells were strengthened by the fact that ISRIB (which reverses inhibitory effects of eIF2α phosphorylation) was unable to restore translation under these stress conditions (chronic Tg + acute SA) (**Figure 3C**), but not when treated with Tg alone for 24h in neuronal cells (**Figure 5B**). Therefore, chronically ER stressed neurons redirect towards an eIF2α-independent mechanism when exposed to oxidative stress. These results are quite unexpected given that GADD34 expression levels are still elevated in these cells (**Figure 3B**), as GADD34 mRNA levels are known to serve as a molecular memory damper to subsequent stresses (Batjargal et al., 2022; Klein et al., 2022; Shelkovnikova et al., 2017). This apparent ability of (at least) glial cells to reset the ISR in the presence of GADD34 while neuronal cells seem to “forget” how to respond brings an important question: was it even meant to be remembered? Given this lack of activation of a subsequent ISR in neuronal cells we consider three possible reasons, but not mutually exclusive, by order of likelihood. (1) The transition to a chronic ISR highlights the inability of neuronal cells to re-shape cell adaptation solely through the ISR, hence shifting towards alternative and/or parallel signalling pathways (*e.g*., mTOR (Guan et al., 2014; Terenzio et al., 2018), eIF2A (Kim et al., 2011), eIF3d (Guan et al., 2017), eEF1A2 (Mendoza et al., 2021). Recent evidence supports that translation repression is maintained in PERK-deficient neurons by complementary eIF2α-independent mechanisms (tRNA-cleaving RNase) (Wolzak et al., 2022). (2) Secondly, a cell non-autonomous trigger of the acute ISR in neuronal cells undergoing chronic stress thus signalled by eIF2-dependent glial cells, supported by recent work were targeting PERK the eIF2α axis of PERK in astrocytes rescues prion-causing neuronal dysfunction (Smith et al., 2020). (3) Thirdly, multiple eIF2α kinases might be activated in neuronal cells (Alvarez-Castelao et al., 2020; Wolzak et al., 2022) during chronic ER stress, thus less susceptible to re-start an acute ISR when subsequently challenged with a different stressor, whereas glial activation of eIF2α kinases may be stimuli-specific. ISR ‘exhaustion’ has also been recently appreciated where translational-demanding cell types (in this study, pancreatic β cells) are susceptible to ATF4-mediated transcriptome decay when faced with frequent ER stress insults (Chen et al., 2022).

Activation of the acute ISR increases eIF2Bδ localisation to small eIF2B bodies (containing γ and ε) with increased eIF2 shuttling in astrocytic cells, suggesting the stress-induced formation and clustering of eIF2Bγδε subcomplexes with increased GEF activity (Hodgson et al., 2019). Here we observed that while eIF2Bδ redistributes to small bodies during acute Tg (1h) and SA (0.5h) treatments it returns to basal levels upon chronic Tg treatment (24h) (**Figure 4A**); thus, suggesting subunit remodelling of small eIF2B bodies is a transient event and specific to the acute ISR independent of cell types, and reversed during the chronic ISR. Interestingly, a subsequent SA treatment to Tg pre-treated cells, which selectively re-elevates eIF2α phosphorylation in U373 and MO3.13 cells (**Figure 3B**), is also accompanied by increased eIF2Bδ localisation of small bodies in these cells but not in SH-SY5Y cells (**Figure 4A**). Surprisingly, we show that eIF2Bδ localisation to small eIF2B bodies in the presence of acute ER stress is only partially dictated by eIF2α phosphorylation (**Figure 4B**), suggesting that non-ISR mechanisms are at play in eIF2B body remodelling. We do not discard the possibility that chemical stressors such as Tg may trigger multiple pathways (Li & Hu, 2015; Wink et al., 2017) that could influence our observations. Given that ER stress-induced eIF2Bδ remodelling exists upon PERK inhibition (yet at a lower level), we speculate that these other pathways could serve as an activator of eIF2B body remodelling, further enhanced and/or maintained by eIF2α phosphorylation.

As shown previously, ISRIB mimicked the acute ISR program response in astrocytic cells (Hodgson et al., 2019) and oligodendrocytic cells, however chronic ER stress hindered the action of ISRIB for the latter (**Figure 5A**). However, this “stress-mimicking” feature of ISRIB was not recapitulated in neuronal cells. ISRIB alone or in combination with chronic ER stress did not increase eIF2Bδ localisation in neuronal small eIF2B bodies (**Figure 5A**). Unexpectedly, despite the differences in eIF2Bδ movement, ISRIB treatment of chronic ER-stressed cells had a unanimous restorative effect on translation across all cell types (**Figure 5B**), thus suggesting that ISRIB-mediated translational rescue does not require increased eIF2Bδ-containing small bodies as a general feature. However, our data suggest that eIF2Bδ regulatory remodelling may be functionally relevant to astrocytic cells.

eIF2B regulatory subunits control eIF2B activity upon stress (Krishnamoorthy et al., 2001; Pavitt et al., 1997) whilst catalytic subunits remain desensitized to stress when uncoupled from regulatory subunits (Liu et al., 2011; Wortham et al., 2014). Indeed, FRAP analysis identified a cell type specific correlation between eIF2B subunit composition and eIF2 recovery upon acute ISR and ISRIB treatment. (1) In neuronal cells, acute ER stress had a predominantly inhibitory effect on the eIF2 recovery of small and large bodies (**Figure 6A**). This may be a result of a more homogenous composition of both small and large bodies hence functionally similar. This agrees with our previous observations that neuronal small bodies have increased regulatory subunit composition (**Figure 1D**). (2) In contrast, astrocytic cells displayed distinctive functional responses for small and large eIF2B bodies as previously reported (Hodgson et al., 2019): small bodies exhibited enhanced eIF2 recovery upon acute ER stress and ISRIB treatment, while acute ER stress repressed the eIF2 recovery of large bodies which is rescued by the addition of ISRIB (**Figure 6A**). (3) Finally, acute ER stress and ISRIB had no impact on small and large eIF2B bodies of oligodendrocytic cells. This unresponsiveness is likely related to the lack of eIF2Bβ from eIF2B bodies (**Figure 1D**), hence supporting that non-decameric eIF2B subcomplexes may predominantly localise to eIF2B bodies in oligodendrocytic cells.

While acute ER stress shows cell type-dependent discrepancies in the % recovery of eIF2 of eIF2B bodies, these changes are unanimously reversed upon sustained Tg treatment (**Figure 6B**), further suggesting that eIF2B localisation is normalised during chronic ISR. Guan *et al*. recently provided evidence that recovery of eIF2B activity may not be required upon transition to chronic stress and may be alternatively mediated via eIF3 (Guan et al., 2017).

A cell-specific relationship between eIF2Bδ redistribution and eIF2 recovery was observed in astrocytic cells as illustrated in Figure 7. Indeed, ISRIB treatment has a dominant effect of increasing eIF2δ composition of small bodies, either alone or in combination with chronic ER stress (**Figure 5A**), accompanied by enhanced % recovery of eIF2 (**Figure 6B**). This relationship is not recapitulated in the other cell types used in this study, which requires further *in vitro* studies to investigate the cell specific GEF activity of eIF2Bγδε subcomplexes.

**Figure 7.**
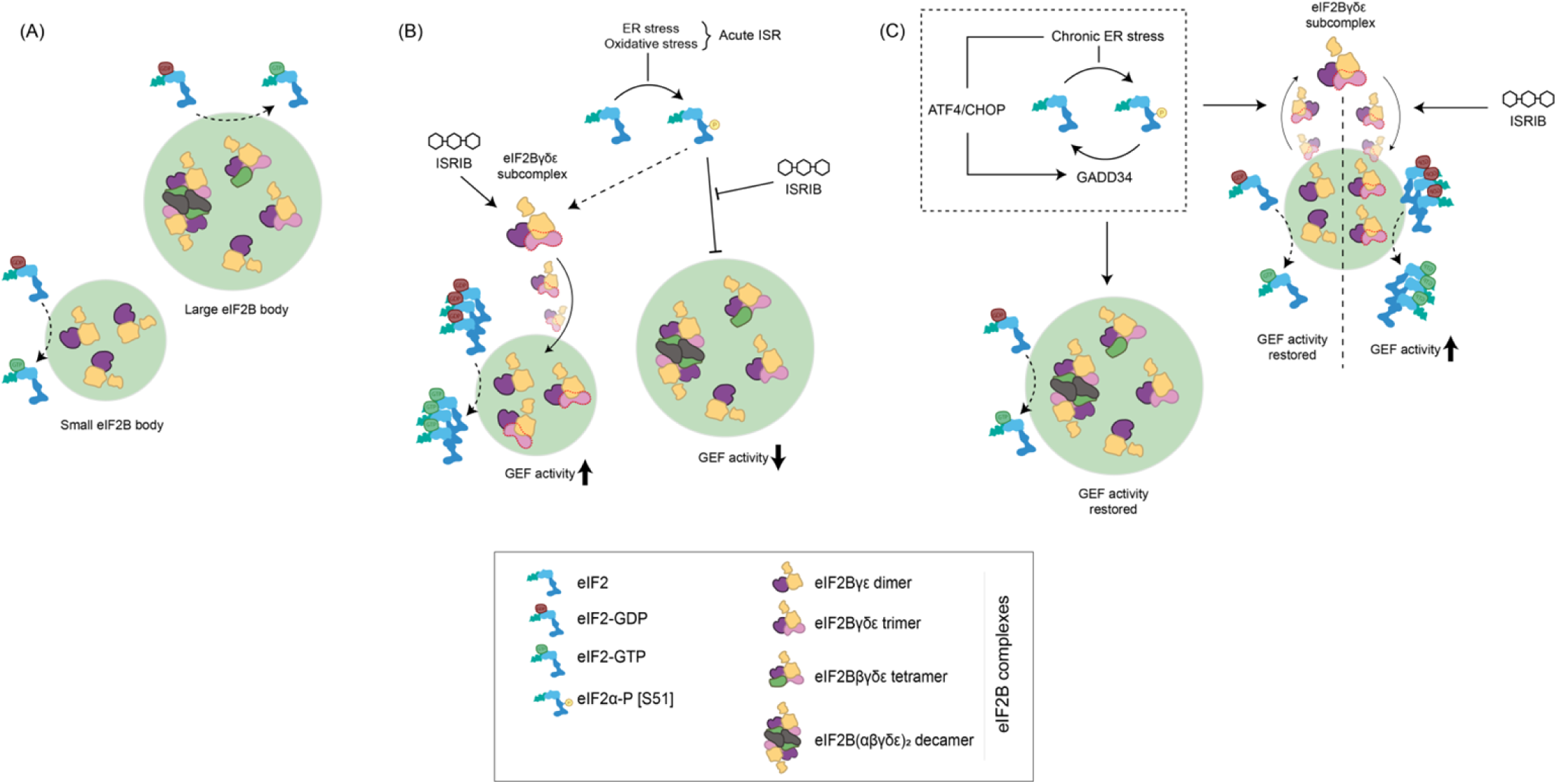
Working model for the impact of cellular stress and ISRIB in eIF2B bodies of astrocytes. **(A)** eIF2B localises to small eIF2B bodies containing catalytic subcomplexes and larger eIF2B bodies containing a variety of regulatory subcomplexes (including decameric eIF2B). **(B)** Upon activation of the acute ISR program, eIF2Bγδε subcomplexes are formed and localised to small eIF2B bodies which we hypothesize to have a regulatory role in eIF2B GEF activity; whilst large eIF2B bodies are negatively impacted. **(C)** During transition to a chronic ISR, eIF2Bδ distribution in small bodies is reversed and GEF activity is restored to basal rates, whereas ISRIB treatment bypasses transient eIF2Bδ distribution by prompting extended eIF2Bγδε formation by direct interaction with eIF2Bδ.

Collectively, our results demonstrate that cells display cell type specific localisation and regulation of eIF2B bodies. The existence of different eIF2B subcomplexes of eIF2B bodies may allow unique rates of ternary complexes levels and adaptability to stress which overall might make translation more efficient and/or more easily regulated. More importantly, we provide evidence for cell type-specific fine-tuning of eIF2B function and regulation, the core event of the ISR; further emphasizing the need to tailor therapeutic interventions in a cell type-manner.

## Acknowledgements

The authors wish to thank Dr Truus Abbink (Amsterdam UMC) for insightful discussions and helpful suggestions. We thank Professor Nicola Woodroofe (Sheffield Hallam University) for kindly gifting us the MO3.13 cells and selective marker antibodies. We gratefully thank the Biomolecular Sciences Research Centre and Sheffield Hallam University for funding this work.

## Statements and Declarations

The authors have no relevant financial or non-financial interests to disclose.

## Material and Methods

### Cell culture

Human U373 astrocytoma cells were cultured in Minimum Essential Medium (MEM), supplemented with 10 % (v/v) heat-inactivated fetal bovine serum (FBS), 1 % (w/v) Non-essential amino acids, 1 % (w/v) sodium pyruvate, 2 mM L-glutamine and 1 % (w/v) penicillin/streptomycin. Human SH-SY5Y dopaminergic neuroblastoma cells were cultured in Dulbecco’s modified Eagle’s medium:F-12 (DMEM:F-12; 1:1) containing high glucose (3.151 g/L) (Lonza), supplemented with 10% (v/v) heat-inactivated fetal bovine serum (FBS), 2 mM L-glutamine and 1% (w/v) penicillin/streptomycin. Human MO3.13 hybrid primary oligodendrocytes were cultured in high glucose Dulbecco’s modified Eagle’s medium (DMEM), supplemented with 10 % (v/v) heat-inactivated foetal bovine serum (FBS) and 2 mM L-glutamine. All experiments were done with passage number no higher than 25. All media and supplements were purchased from Life Technologies Co. (New York, USA), except when indicated otherwise. All cell lines were maintained at 37°C under 5% CO2 and were routinely tested for contamination with MycoAlert^™^ Mycoplasma Detection Kit (Lonza).

### Cell treatments

For acute/transient induction of the ISR, cells were treated with 1 μM thapsigargin (Tg) (Sigma-Aldrich) for 60 minutes; 3 μg/mL tunicamycin (Tm) (Cayman Chemical) for 2h; and 125 μM sodium arsenite (SA) (Sigma-Aldrich) for 30 minutes. For chronic induction of the ISR, cells were treated with 300 nM Tg for 24h. For acute/transient cellular stress previously challenged with a chronic induction of the ISR, cells were treated with 300 nM Tg for 24h where 1 μM Tg, 3 μg/mL Tm or 125 μM SA were added in the last 1h, 2h and 30 minutes, respectively. For ISRIB treatment, cells were added with 200 nM ISRIB (Sigma, Dorset, UK) for 1h. For PERK inhibition treatment, cells were treated with 500 nM GSK2606414 (#5107, Tocris) for 1h. As control, cells were treated with vehicle solution (DMSO) with the highest volume and treatment duration depending on its respective experimental setup.

### Plasmids

pCMV6-AC-tGFP plasmid vector encoding EIF2B5 (#RG202322) and EIF2S1 (#RG200368) was purchased from OriGene (Rockville, Maryland, USA). The coding open reading frame of EIF2B5 from the pCMV6-AC-tGFP vector was cloned into an empty pCMV6-AC-mGFP (#PS100040, OriGene) and empty pCMV6-AC-mRFP (#PS100034, OriGene) vectors. The constructs were verified by sequencing.

### Transient transfection procedures

U373, SH-SY5Y and MO3.13 cells were seeded at a density of 3×10^5^, 5×10^5^ and 2.5×10^5^ cells, respectively, in a 6-well plate for at least 24 hours before transfection. For U373 cells, transient transfection was performed with transfection reagent 25-kDa polyethylenimine, branched (PEI) (1 mg/mL) (Sigma-Aldrich). 1 μg of plasmid DNA was diluted in 100 μL of serum- and antibiotic-free medium and incubated with 4 μg PEI for 10 minutes. 600 μL of antibiotic-free media was added to the transfection mixture, added to cells, and incubated for 4 hours at 37°C. Media was removed and replaced with complete media and incubated for 24-48h at 37°C prior to confocal imaging. SH-SY5Y and MO3.13 cells were transfected with Lipofectamine 3000 according to manufactures instructions.

### Immunoblotting

5×10^5^ cells were cultured on 6-well plates. Whole-cell protein lysates were prepared in CelLytic M lysis buffer (Sigma) with 1% protease/phosphatase inhibitors (Sigma) and other supplements (17.5 mM β-glycerophosphate, 1 mM PMSF, 10 mM NaF). Lysates were incubated on ice for 10 minutes and centrifuged (13,000 rpm, 10 min, 4°C) to remove cellular debris. Protein concentrations were determined with Qubit™ Protein Assay Kit (Thermo-Fisher) and subjected to SDS-PAGE electrophoresis. For western blots, samples were run on 7.5 or 10 % polyacrylamide gel and transferred using Trans-Blot Turbo Mini-nitrocellulose Transfer packs (Bio-Rad) on a Trans-Blot Turbo apparatus. When necessary, membranes were subjected to Revert^™^ Total Protein Stain (LiCor) following manufacturer’s instructions. Membranes were blocked in 5 % milk or 5 % BSA and probed with primary antibodies diluted in 5 % milk or 5 % BSA, overnight at 4°C. The following antibodies were used: eIF2α (1:500, ab5369, Abcam), phosho-eIF2α[ser51] (1:500, ab32157, Abcam), PERK (1:1000, 20582-1-AP, Proteintech), GADD34 (1:500, 10449-1-AP, Proteintech), CHOP (1:1000, 15204-1-AP, Proteintech), ATF4 (1:750, ab184909, Abcam), GAPDH (1:5,000, #2118, Cell Signalling). Membranes were then washed 3 times for 5 minutes/each in TBST, followed by probing with secondary antibodies diluted in 5 % milk or 5 % BSA in TBST for 1h at RT: goat-anti-rabbit IRDye 680RD (1:10,000, 925-68071, LiCor) and goat-anti-mouse IRDye 800CW (1:10,000, 925-32210, LiCor) and washed 3 times for 5 minutes/each in TBST. Membranes were visualised and quantified on a LiCor Odyssey Scanner with Image Studio Lite software.

### Puromycin incorporation assay

For puromycin (PMY) integration, 91 μM puromycin dihydrochloride (Thermo Fisher Scientific) was added to media 5 minutes prior to harvesting and incubated at 37°C. Cells were washed twice with ice-cold PBS supplemented with 355μM cycloheximide (Calbiochem), lysed and immunoblotted as described previously. Primary PMY-specific antibody (1:500, clone 12D10, MABE343, Sigma-Aldrich) was used to detect puromycinylated proteins. GAPDH was used as loading control.

### Immunocytochemistry

22×22 mm squared glass coverslips (Sigma-Aldrich) were rinsed with 100% IMS (Fisher Scientific), added to 6-well plates, and left until IMS fully evaporated. Cells were seeded and transfected as described previously. U373 and SH-SY5Y cells were fixed in ice-cold methanol (Fisher Scientific) at −20 °C for 15 min, and MO3.13 cells in 4 % (w/v) paraformaldehyde (Alfa Aesar) at RT for 20 minutes. For methanol fixation, cell membranes were washed with PBS supplemented with 0.5 % (v/v) Tween 20 (PBST), 3 times for 3 minutes and then blocked in PBS supplemented with 1% (w/v) bovine serum albumin (BSA) for 60 minutes at RT, or overnight at 4°C, under gentle shaker. For paraformaldehyde fixation, cells were washed 3 times with PBST for 3 minutes, permeabilized with 0.1 % (v/v) X-Triton for 5 minutes at RT and blocked in 1 % (w/v) BSA in PBST for 60 minutes at RT or overnight at 4°C, under gentle shaker. Cell membranes were probed with primary antibodies diluted in 1 % (w/v) BSA in PBS, overnight at 4°C under gentle shaker, as following: eIF2Bα (1:25, 18010-1-688 AP, Proteintech), eIF2Bβ (1:25, 11034-1-AP, Proteintech), eIF2Bγ (1:50, sc-137248, Santa Cruz), eIF2Bδ (1:50, sc-271332, Santa Cruz), eIF2Bε (1:25, HPA069303, Sigma-Aldrich). Cells were then washed 3 times with PBST for 5 minutes, followed by probing with the appropriate host species Alexa Fluor conjugated secondary antibody (Thermo Fisher Scientific), diluted in 1 % (w/v) BSA in PBS, for 60 minutes at RT. Following secondary antibody incubation, cells were washed with PBST, 3 times for 5 minutes, and mounted with ProLong^™^ Gold Antifade Mountant with DAPI (Invitrogen, Thermo Fisher Scientific). Cells were visualised on a Zeiss LSM 800 confocal microscope.

### Confocal imaging and fluorescence recovery after photobleaching (FRAP)

Imaging was performed using a Zeiss LSM 800 confocal microscope combined with Zeiss ZEN 2.3 (blue edition) software for data processing and analysis. Both 40x and 63x plan-apochromat oil objectives were used and a laser with maximum output of 10mW at 0.2 % (488 nm) and 5.0 % (561 nm) laser transmissions. Fluorescence crosstalk was minimal and bleed-through was not observed. Image acquisition was performed by orthogonal projection of a Z-stack of automatically calculated increments for complete single cell imaging. FRAP analysis was performed to quantify the shuttling rate of eIF2 through localised eIF2B as described in the methodology by (Hodgson et al., 2019). FRAP experiments were carried out by live cell imaging in an incubation chamber with appropriate temperature and CO_2_ levels. Specific areas containing an entire cytoplasmic eIF2α-tGFP foci were manually marked for bleaching using 23 iterations at 100 % laser transmission (488 nm argon laser). Pre-bleaching image and intensity of targeted foci (ROI – region of interest) was captured followed by 44 images captured every 151 ms for a total of 7.088 s. In-cell fluorescence intensity was captured to normalise against ROI. Out-of-cell fluorescence, or background intensity (B), was measured and subtracted from ROI and T values to provide corrected measurements. Normalised data was fitted to a one-phase association curve using GraphPad Prism to quantify rate of recovery. The relative percentage of eIF2 recovery was determined as the plateau of the normalised FRAP curve.

### Statistical analysis

All statistical assessments were made in GraphPad Prism 7 software, with a significance at p<0.05. All data is presented as means ± standard errors of the mean (s.e.m.). Data was subjected to Shapiro-Wilk normality test. If parametric, data was analysed by one-way ANOVA test for comparison of three of more groups followed by Tukey’s correction post-hoc test. If non-parametric, data was analysed by Kruskal-Wallis test for comparison of three of more groups followed by Dunn’s correction post-hoc test.

---

**Figure S1.**
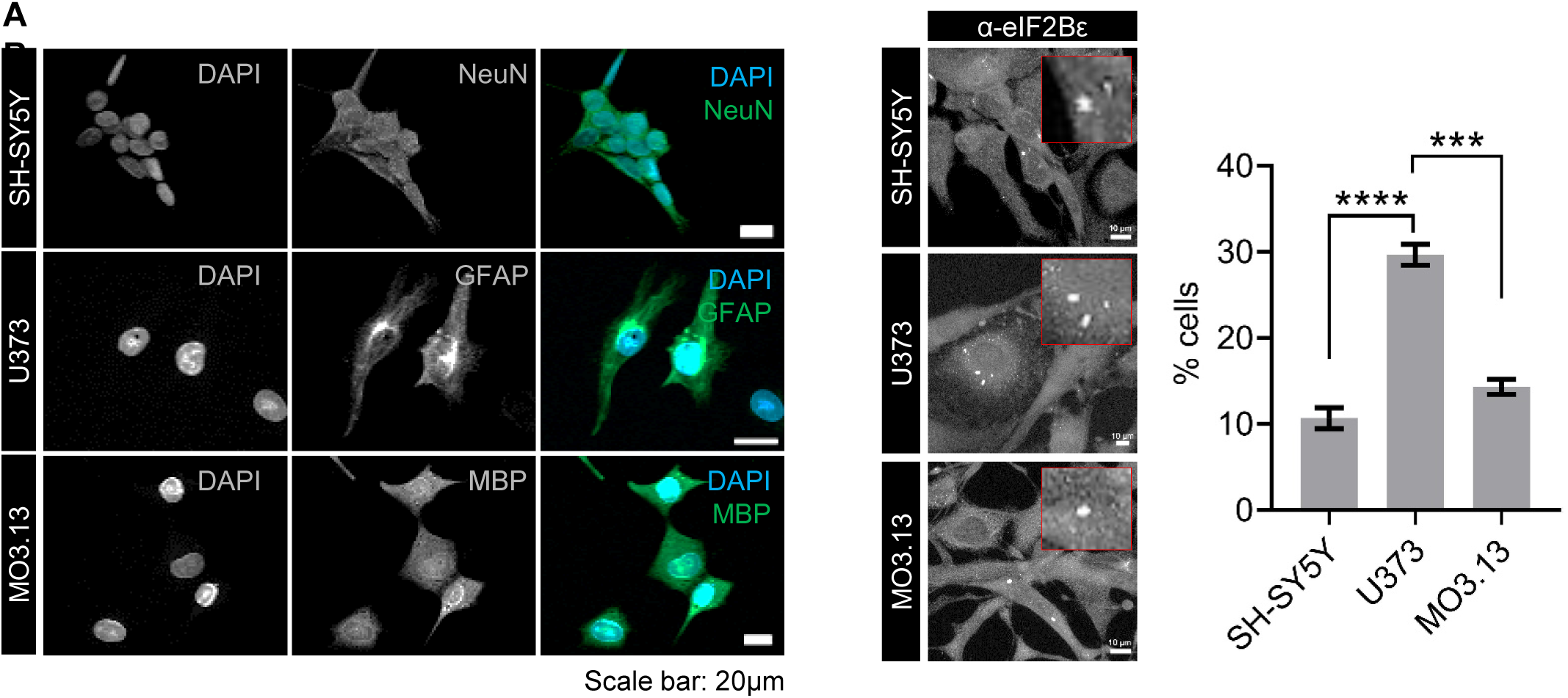
Endogenous eIF2B bodies are expressed in SH-SY5Y, U373 and MO3.13 cells. **(A)** Confocal images of SH-SY5Y, U373 and MO3.13 cells immunostained for neural marker *NeuN*, astrocytic marker *GFAP* and oligodendrocytic marker *MBP*, respectively. **(B)** Confocal images of SH-SY5Y, U373 and MO3.13 cells immunostained with α-eIF2Bε (*left panel*). Mean percentage of cells displaying at least one eIF2Bε-containg body in a population of 100 cells (mean±SEM; *N*=3; \*\*\*\**p*≤0.0001, \*\*\**p*≤0.001 according to one-way ANOVA) (*right panel*). Scale bar: 10 μm.

**Figure S2.**
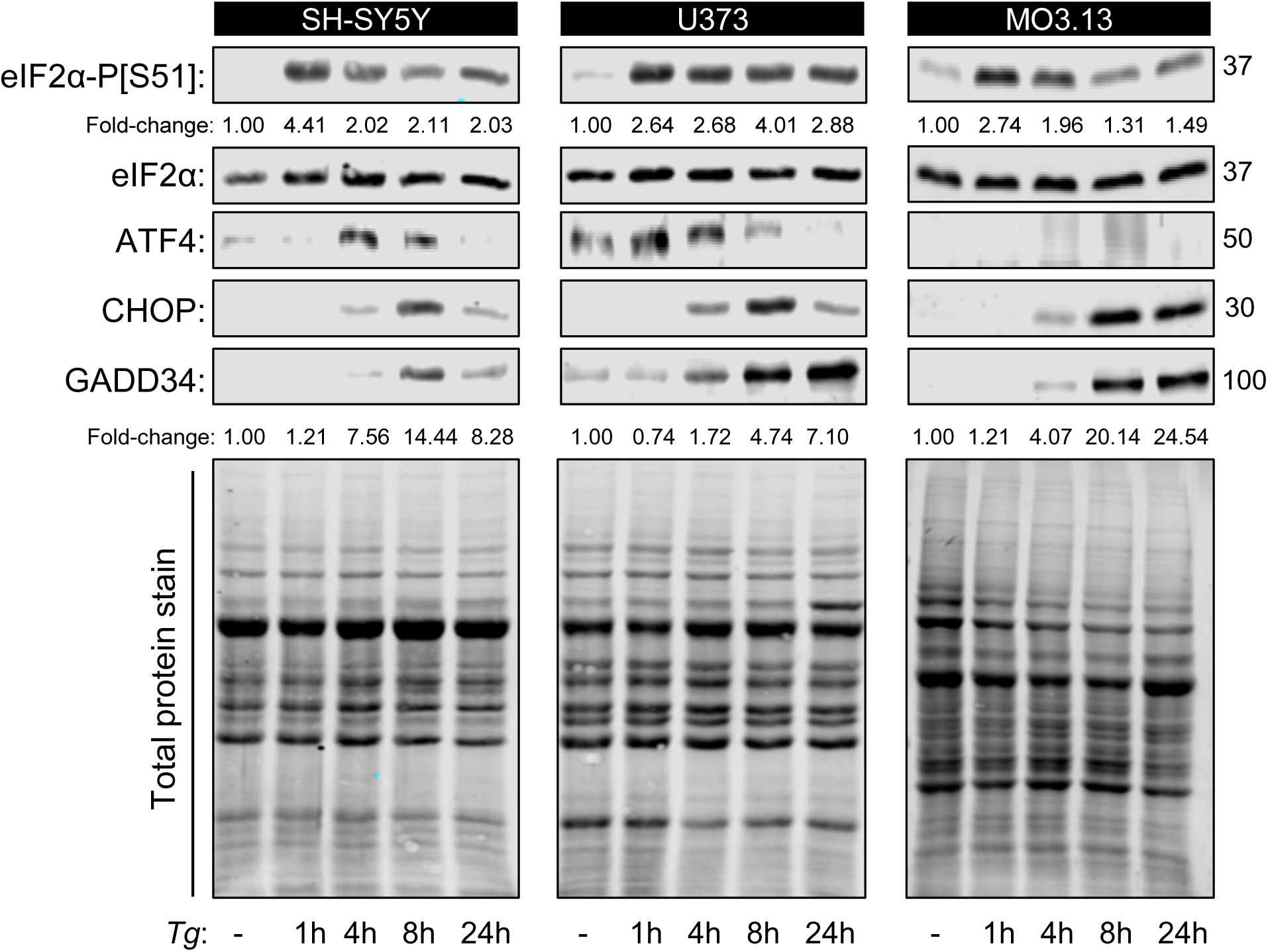
ATF4 expression peaks before 24h time point of Tg exposure. Representative western blot of the ISR expression profile (eIF2α-P[S51], pan-eIF2α, ATF4, CHOP and GADD34) and total protein staining (loading control) in SH-SY5Y, U373 and MO3.13 cells treated with vehicle (DMSO) for 24h or thapsigargin for 1, 4, 8 and 24 hours.

**Figure S3.**
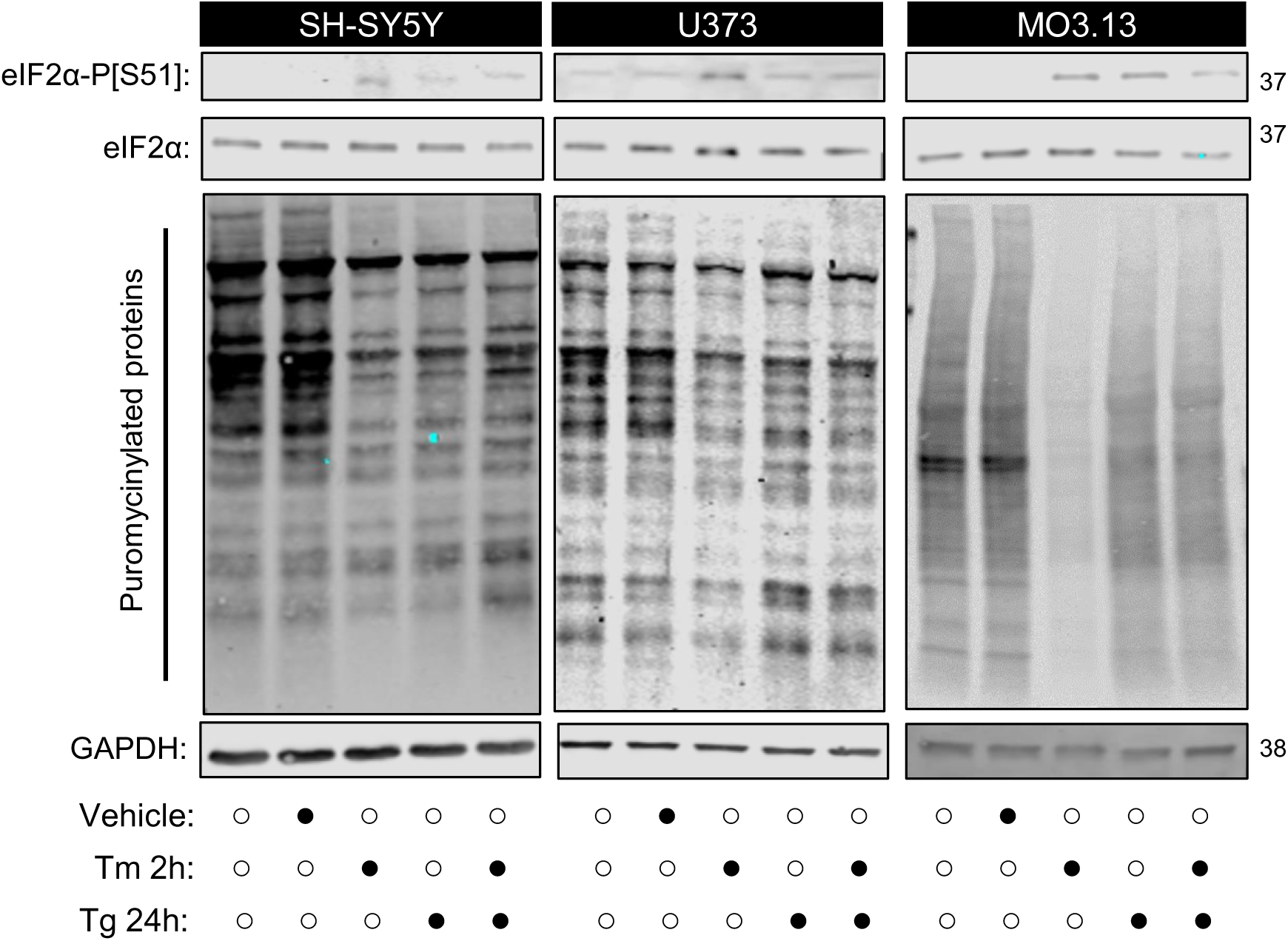
Tunicamycin treatment prior Tg preconditioning for 24h does not further impact on eIF2α-P[S51] levels and protein synthesis. Western blot of the ISR expression profile (eIF2α-P[S51], pan-eIF2α) and global newly synthesized proteins (puromycin incorporation assay) in SH-SY5Y, U373 and MO3.13 cells treated with vehicle (DMSO), acute ER stress inducer (Tm 3 μg/mL for 2h), chronic ER stress (Tg 300nM for 24h) and chronic ER stress further challenged with acute ER stress inducer (Tg 300nM for 24h + Tm 3 μg/mL on the last 2h).

**Figure S4.**
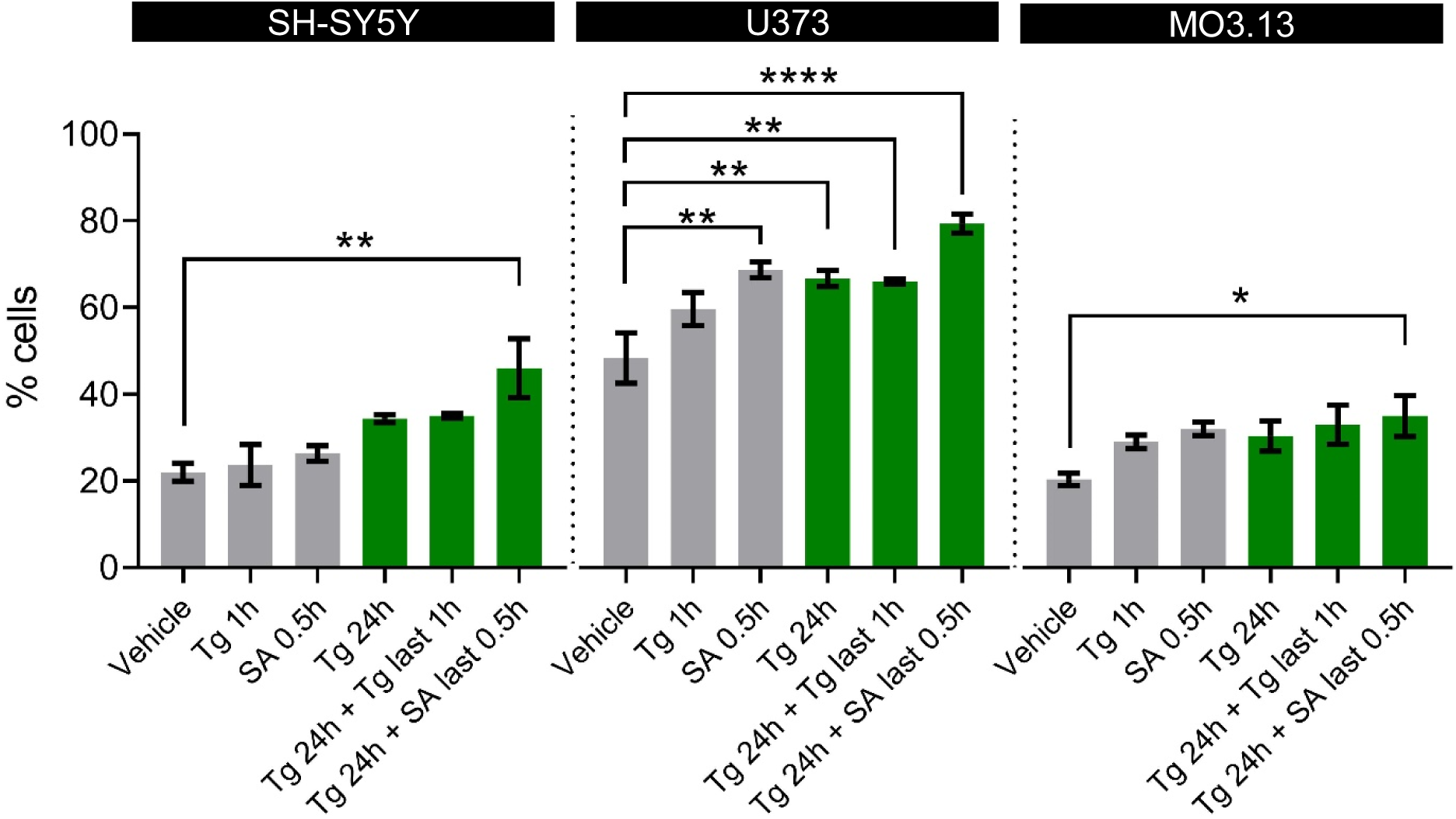
eIF2B localisation increases with cellular stress. SH-SY5Y, U373 and MO3.13 cells were subjected to acute stress inducers (Tg 1μM for 1h and SA 125μM for 30min); chronic ER stress (Tg 300nM for 24h) or subsequently challenged with previously described acute stress treatments (Tg 24h + Tg last 1h; Tg 24h + SA last 0.5h). Mean percentage of cells displaying localised eIF2B (at least one eIF2B body present) in a population of 100 transfected cells (mean±SEM; *N*=3).

## Notes

### Competing Interest Statement

The authors have declared no competing interest.

### Summary of Updates

Authors addresses have been updated

